# Overlapping Representation of Basic Tastes in the Human Gustatory Cortex

**DOI:** 10.1101/2021.10.31.466657

**Authors:** Du Zhang, Xiaoxiao Wang, Yanming Wang, Benedictor Alexander Nguchu, Zhoufang Jiang, Chenwei Shi, Bensheng Qiu

## Abstract

The topological representation is a fundamental property of human primary sensory cortices. The human gustatory cortex (GC) responds to the five basic tastes: bitter, salty, sweet, umami, and sour. However, the topological representation of the human gustatory cortex remains controversial. Through functional magnetic resonance imaging(fMRI) measurements of human responses to the five basic tastes, the current study aimed to delineate the taste representations within the GC. During the scanning, the volunteers tasted solutions of the five basic tastes, then washed their mouths with the tasteless solution. The solutions were then sucked from the volunteers’ mouths, eliminating the action of swallowing. The results showed that the bilateral mid-insula activated most during the taste task, and the active areas were mainly in the precentral and central insular sulcus. However, the regions responding to the five basic tastes are substantially overlapped, and the analysis of contrasts between each taste response and the averaged response to the remaining tastes does not report any significant results. Furthermore, in the gustatory insular cortex, the multivariate pattern analysis (MVPA) was unable to distinguish the activation patterns of the basic tastes, suggesting the possibility of weakly clustered distribution of the taste-preference neural activities in the human insular cortex. In conclusion, the presented results suggest overlapping representations of the basic tastes in the human gustatory insular cortex.

## 1 Introduction

Much of the mammal primary sensory, such as somatosensory, visual and auditory cortex, is organized into topological maps: nearby neurons have similar sensory receptive fields (Romani, Williamson, & Kaufman, 1982; Sanchez-Panchuelo, Francis, Bowtell, & Schluppeck, 2010; Wandell, Dumoulin, & Brewer, 2007). Taste quality is generally acknowledged to contain five perceptual dimensions -- bitter, salty, sour, sweet and umami (McBurney & Gent, 1979; Simon, de Araujo, Gutierrez, & Nicolelis, 2006). For taste, the primary gustatory cortex was located in the middle insula by previous works (Avery et al., 2020; Small, 2010; Veldhuizen et al., 2011; Yeung, Goto, & Leung, 2018). However, the debate about its topography within the gustatory cortex lasts for years. X. Chen, Gabitto, Peng, Ryba, and Zuker (2011) observed highly separated clusters of neurons responding to taste qualities in the mouse. In controversial, more recent researches observed overlapped clusters of taste-responsive neurons in the mouse GC (K. Chen, Kogan, & Fontanini, 2021; Fletcher, Ogg, Lu, Ogg, & Boughter, 2017). Most taste-responsive neurons within the macaque insular cortex seemed to tune to multiple tastes, with no clear topographic organization of taste sensitivity (Scott & Plata-Salaman, 1999). Many human functional magnetic resonance imaging (fMRI) studies explored the question of a gustotopic organization of the insular cortex. Some of them failed to find spatial segregation of taste qualities (Dalenberg, Hoogeveen, Renken, Langers, & ter Horst, 2015; Schoenfeld et al., 2004). Some reported gustotopic organization but varied across participants (Prinster et al., 2017). Over the past few years, the 7Tesla high-field MRI offered the opportunity of investigating human brain activity in a higher signal-to-noise ratio (SNR) and higher spatial resolution. The 7Tesla fMRI studies also failed to find the brain regions that consistently preferred any individual taste (Avery et al., 2020; Chikazoe, Lee, Kriegeskorte, & Anderson, 2019).

The results of these gustatory fMRI may be affected by some nuisance effects. The head motion is one of the most significant confoundings and fMRI is extreme sensitivity to head motion (Matthews & Jezzard, 2004). Swallowing would cause inescapability task-related head motion and induce related somatosensory (S. L. Yang, T. J. Ross, Y. Q. Zhang, E. A. Stein, & Y. H. Yang, 2005). The temperature of the tastants could also contribute to the variability of blood oxygen level-dependent (BOLD) response in the insular cortex (Guest et al., 2007). Thus, based on Canna et al. (2019)’s low-cost taste delivery equipment, we built a gustometer equipment furtherly including a pump for sucking liquid from the subject’s mouth and water bathing to maintain the tastants’ temperature. It is also possible that the location of the gustotopic map might vary across subjects, and it would mess up the results of group analysis. So extensive fMRI measurements (180 trials per taste per subject) were conducted by our self-made gustometer and novel taste task design, strictly controlling for the head movement. Both univariate analyses and multivariate analysis were employed to explore the taste coding in the human insular cortex.

## 2 Materials and Methods

### 2.1 Participants

Four subjects (2 female) aged between 21 and 26 (average: 24.5years) participated in the taste fMRI study. All participants were right-handed, and ethics approval for this study was granted by the University of Science and Technology of China. Participants were excluded from this experiment if they had any history of prenatal drug exposure, current usage of psychotropic medications, neurological illness, or any exclusion criteria for magnetic resonance imaging (MRI). The participants were asked to abstain from food for two hours before participating in experiments.

### 2.2 Taste solution

Five tastant solutions were administered to our participants during scanning, namely: bitter (0.5mM quinine hydrochloride), salty (180mM NaCl), sour (10mM citric acid), sweet (2mM sucralose), umami (100mM Monosodium Glutamate + 5mM Inosine monophosphate). The tasteless baseline solution was composed of 12.5mM KCL + 1.25mM NaHCO3. The choice of concentrations for each taste referred to previous neuroimaging studies of taste perception (Han, Mohebbi, Unrath, Hummel, & Hummel, 2018; Mascioli et al., 2015; Prinster et al., 2017). Every tastant solution was dissolved in the purified water. The tasteless solution was used as the control to remove the confounding effects of other feelings caused by introducing a fluid into the mouth.

### 2.3 Gustometer description

A self-made MRI-compatible system was designed to deliver tastants through the peristaltic pump. The uniformity of the temperature of tastant solutions was maintained using a water bath. A single-chip microcontroller STC12C5A60S2 was used to control the supply of the solutions to the subject. The solutions were flowing through the peristaltic pumps and delivered chronologically (Fig.1A). These peristaltic pumps are software-driven, capable of managing the flow rate, being set at 1.27ml/s in our case. Deliverance and removal of tastant solutions were achieved by a well-fabricated tooth socket (Fig.1B). The device was inserted correctly into the subject’s mouth during experiment setup to ensure that participants need not swallow during taste fMRI. Visual cues for fMRI tasks were projected onto a screen located inside the scanner bore and viewed through a mirror system mounted on the head-coil. Tastant delivery and visual stimulus presentation were controlled and synchronized via a custom-built program developed in the Psychtoolbox-3 (Brainard, 1997; Kleiner, Brainard, & Pelli, 2007; Pelli, 1997) and Matlab (Mathworks, Natick, MA) environment.

**Figure 1.**
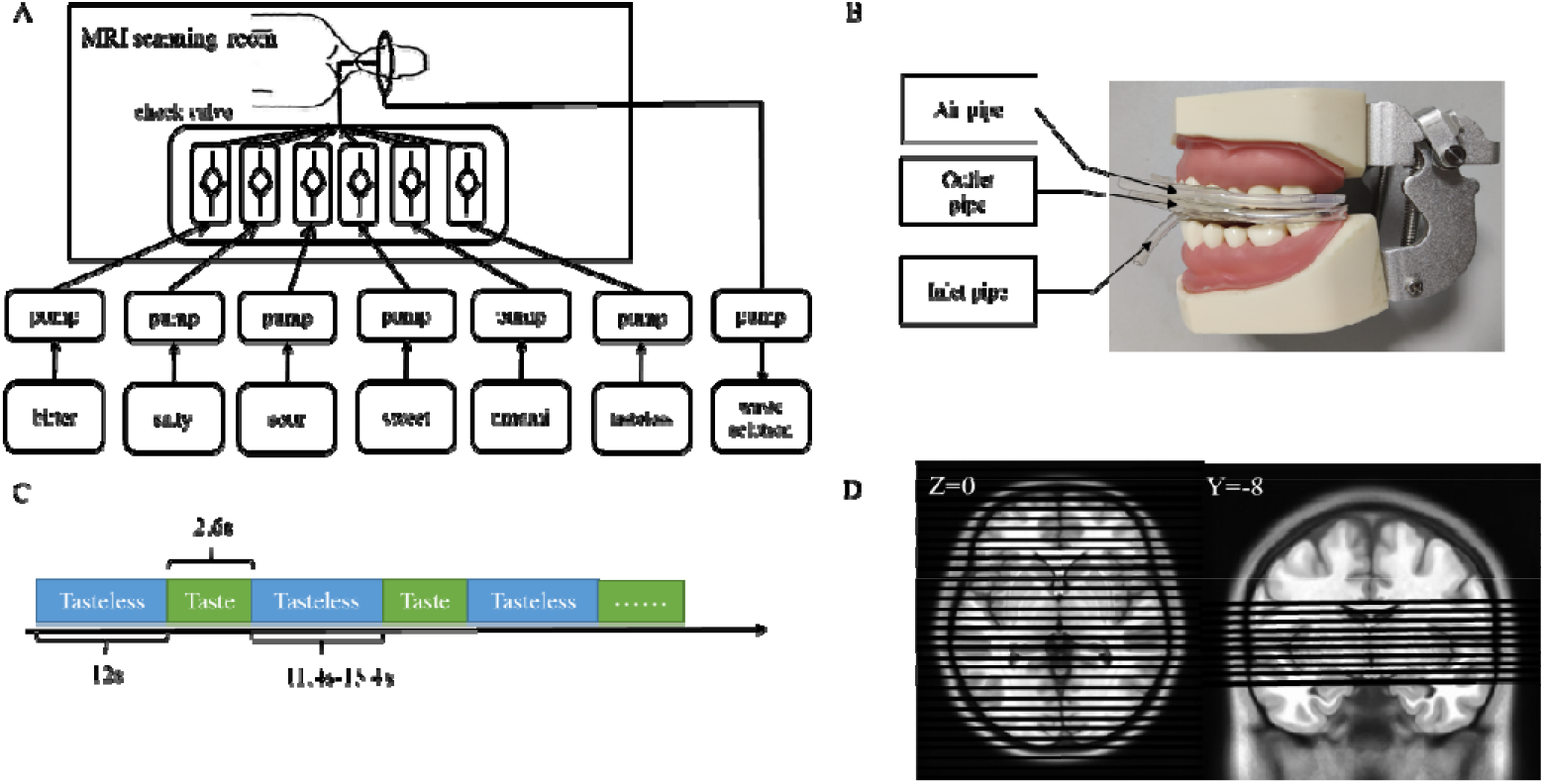
Gustometer and taste fMRI design description. **A**. The taste solution delivery system. Six tastant solutions were delivered to the homemade tooth socket in the subjects’ mouth by peristaltic pumps controlled by a single-chip microcontroller. The flow rate was controlled at 1.27 ml/s. Check valves ensure one-way liquid flow, and an independent peristaltic pump extracted the waste solution during the taste fMRI task. **B**. The homemade tooth socket. The six solution-inlet tubes were connected to the anterior inlet pipe and transferred the taste solution to the tip of the tongue. The outlet pipes were attached at the posterior side of the tooth socket and extracted the solution from the tongue base so that subjects were unnecessary to swallow the fluids. **C**. The diagram of the taste task in fMRI. A tasteless stimulus begins, and a queue of 30 trials whose average length was 16s with a taste stimulation for 2.6s and a tasteless inspiration for 11.4-15.4s. **D**. Image acquisition. An axial (left) and coronal (right) view of the image-acquisition areas of the brain. Actual scanning contained 30 slices.

### 2.4 Behavioral tests

Before the fMRI taste perception task, all subjects underwent behavioral tests to ensure that everyone could discriminate the tastes and correctly identify the solutions. The subjects rated each taste solution’s perceived intensity and pleasantness using a visual analog scale from 0 for very weak to 10 for very intense, which was already used and validated in the previous study(Prinster et al., 2017). A paired t-test was used to analyze the behavioral data, with a significance threshold of P<0.05.

### 2.5 Taste task in fMRI

The taste task begins with a tasteless for 12s, then follows a queue of 30 trials with an average length of 16s with a taste stimulation for 2.6s and a tasteless inspiration for 11.4-15.4s (Fig. 1C). For maintaining alarm, the subjects were asked to press the response button through their right thumb once they felt the taste. The sequence of tastes was pseudo-randomized, in which each taste was repeated six times, and the tastes of neighboring trials were different during each scan. Each scan lasted for 492 seconds (8min12sec). The scan was repeated six times per session in five sessions, with one session per day. Thus, we obtained 180 trials of fMRI scan for each of the five tastes of each participant.

### 2.6 FMRI data Acquisition

MRI data scanning was performed at the MRI center of the medical center of the University of Science and Technology of China. MRI images were acquired with the aid of a 3-T scanner (Discovery 750; GE Healthcare, Milwaukee, WI) and an 8-channel head coil. T1-weighted images were obtained to provide an anatomical reference for the data analysis, with repetition time (TR) = 8.2ms, echo time (TE) = 3.2 ms, flip angle =12°, and voxel size =1×1×1mm^3^. The human GC was found to be in the insular cortex. To get a relatively high spatial resolution in the limited acquisition time, the field of view (FOV) of the taste task fMRI covered partial brain, i.e., the whole insular cortex and nearby brain areas(Fig. 1D), with 30 axial slices, TR = 2000ms, echo time (TE) = 30ms, flip angle =90°, and voxel size =2×2×2mm^3^. A whole-brain fMRI scan with the same parameter (TR = 2000ms, TE = 30ms, flip angle =90°, voxel size = 2×2×2mm^3^, NEX = 5) was also obtained as registration reference for the taste task fMRI in the following pre-processing steps.

### 2.7 FMRI data analysis

#### 2.7.1 Univariate analysis

Taste fMRI data analysis was performed using the AFNI software (Cox, 1996). The first five volumes of each run were removed. Functional images were then slice timing corrected and realigned within each session. Each TR’s head movement relative to the previous TR was calculated, and a threshold of outlier fraction of 0.2 was used for censoring in the following regression analysis. One run of fMRI data of the Subject 3 was excluded from the data because of excessive head motion (the mean outlier fraction of 28^th^ to 39^th^ TR was 0.424). Both the taste task fMRI volumes and structural MRI were registered to the reference whole-brain fMRI volume. Spatial smoothing with a 2mm full width at half maximum (FWHM) of Gaussian was used.

A General Linear Model was created to estimate the BOLD response to the different conditions. The tasteless solution was the baseline solution whereby the activeness of the brain region was directly determined by the statistical contrast, taste (either bitter, salty, sour, sweet, or umami) versus tasteless. In this model, five regressors of interest for taste (bitter, salty, sour, sweet and umami) trials were made by the convolution of the trial events and the AFNI’s SPMG1 canonical hemodynamic response function. Movement regressors (3 translations, 3 rotations) were also added. A high-pass filter was performed to remove slow-varying signal drifts less than 0.01Hz from the fMRI time series. Within-voxel auto-correlations of time-series data were controlled with AFNI’s 3dREMLfit. Finally, using the ANTs program, the statistic maps were normalized to ICBM152 2009c space through the nonlinear warp from the registered structural data to the 1.0mm isotropic ICBM152 2009c template(Avants, Epstein, Grossman, & Gee, 2008).

At the subject level, to identify those voxels associated with five basic tastes across five sessions, AFNI’s 3dMEMA was utilized with the mating t-statistics and regression coefficients as input. A one-tailed test was used to check for positive effects in five tastes responses. The statistic results were thresholded using a voxel-wise threshold of P<0.01 after the family-wise error (FWE) multiple comparisons, while the Subject3’s threshold was set at P<0.02 due to his/her relatively weak taste reaction. In addition, the statistical mean contrast:(0.2bitter + 0.2salty + 0.2sour + 0.2sweet + 0.2umami) was utilized to corroborate the tastes responsive areas of four subjects and the mean contrast results were FWE corrected thresholding at P<0.005. At the group level, taking within-subject variability into account because there are effect estimates from multiple sessions, regression coefficients for each session were entered into a mixed-effects ANOVA (3dLME in AFNI, z-stats reported) with the session as a random factor (Kaskan, Dean, Nicholas, Mitz, & Murray, 2019). First, we used the statistical contrast: (0.2bitter + 0.2salty + 0.2sour + 0.2sweet + 0.2umami) to identify taste-responsive areas, and the statistical map was corrected for multiple comparisons using the False Discovery Rate (FDR) algorithm (p<0.00001). Second, to identify the brain active areas for five basic tastes, we created group-level maps (bitter, salty, sour, sweet, umami), corrected for multiple comparisons using the False Discovery Rate correction (FDR) algorithm to acquire an overall alpha level of 0.01. We made five taste-responsive areas masks and added them up in a new probability template. Finally, we used the contrast between one taste minus the average responses of other tastes (e.g., bitter - 0.25 salty - 0.25 sour - 0.25 sweet −0.25 umami) to determine brain voxels showing specific taste preferences (3dLME in AFNI; z-stats reported).

To visualize the statistical maps of five taste responses in the insular cortex, freesurfer software (http://surfer.nmr.mgh.harvard.edu/) and Connectome Workbench (https://www.humanconnectome.org/software/connectome-workbench/) were used after multiple comparisons corrected in the volume space. The anatomical surface meshes of the MNI template were reconstructed using Freesurfer’s recon-all program, and the five taste statistical map in MNI space were projected to the cortical surface using FreeSurfer’s mri_vol2surf command.

#### 2.7.2 Multivariate pattern analyses (MVPA)

MVPA was conducted to assess the difference between five basic taste representations of the GC by The Decoding Toolbox (Hebart, Gorgen, & Haynes, 2014). The classification algorithm used for this analysis was linear support-vector-machine (SVM). We modeled each run of taste fMRI tasks separately, leading to 20 beta-coefficient maps for each taste of every subject. Then we trained and tested on subject-level regression coefficient maps using leave-one-out cross-validation (LOOCV). The mid-insula regions-of-interest (ROIs) were identified by mean contrast of five tastes (p<0.00001, FDR corrected for multiple comparisons), as described in univariate analysis.

We also used searchlight (three-voxel radius, including 93 voxels) MVPA with linear SVM kernel to analyze taste responses within the insular cortex. We employed LOOCV to evaluate the activity patterns of different taste types in the searchlight spheres. The output of this searchlight analysis was a map of average classification accuracy minus chance (chanced level = 20%). Then we performed the AFNI program 3dttest++ to make group-level analyses to evaluate the classification results at the group level and performed FDR corrected for multiple comparisons.

## 3 Results

### 3.1 Behavioral results

The range of the taste intensity (*mean*±*St*.*d*.) was 4.75±0.96 for bitter, 7.00±0.82 for salty, 4.50±1.00 for sour, 7.75±0.96 for sweet, 8.80±0.58 for umami, and 2.25±0.50 for neutral. As shown in Table 1, all tastes demonstrated significant differences with the tasteless baseline solution in the intensity rating (P<0.05, paired t-test). Participants enjoyed the sweet taste most (8.25±0.95), followed by umami (5.50±1.91), neutral (5.25±0.50), and salty (4.75±0.96), with bitter (4.25±0.50) and sour (4.50±1.29) being mostly disliked.

**Table1.** Rating results of the intensity and enjoyment of each taste.

| Taste | Rating of the intensity |  | Rating of the enjoyment |  |
| --- | --- | --- | --- | --- |
|  | Mean (St.d.) | vs. tasteless p-value | Mean (St.d.) | vs. tasteless p-value |
| bitter | 4.75 (0.96) | 0.030 | 4.25 (0.50) | 0.092 |
| salty | 7.00 (0.82) | 0.002 | 4.75 (0.96) | 0.391 |
| sour | 4.50 (1.00) | 0.037 | 4.50 (1.29) | 0.391 |
| sweet | 7.75 (0.96) | 0.002 | 8.25 (0.95) | 0.005 |
| umami | 8.50 (0.58) | 0.001 | 5.50 (1.91) | 0.761 |
| tasteless | 2.25 (0.50) |  | 5.25 (0.50) |  |

### 3.2 Brain activations by tastes

#### 3.2.1 Gustatory activation areas with fMRI

We used the mean contrast of BOLD responses to five basic tastes for identifying taste-responsive areas. The gustatory activation areas in the group analysis are shown in Fig.2 and Table 2. The bilateral dorsal mid-insula activated most in the taste-responsive region of the brain. We also observed significant activation of the bilateral dorsal anterior insula, rolandic operculum, tongue somatosensory cortex, the right ventral thalamus, and left cerebellum of lobule VI, in agreement with previous human taste neuroimaging studies(Avery et al., 2020; Small, 2010; Yeung et al., 2018). The rolandic operculum has been identified as a cortical region integrating somatosensory and gustatory information from the tongue (Cerf-Ducastel, Ven de Moortele, MacLeod, Le Bihan, & Faurion, 2001; Mascioli et al., 2015). Our results were obtained using FDR corrected for multiple comparisons (P<0.00001).

**Table 2.**
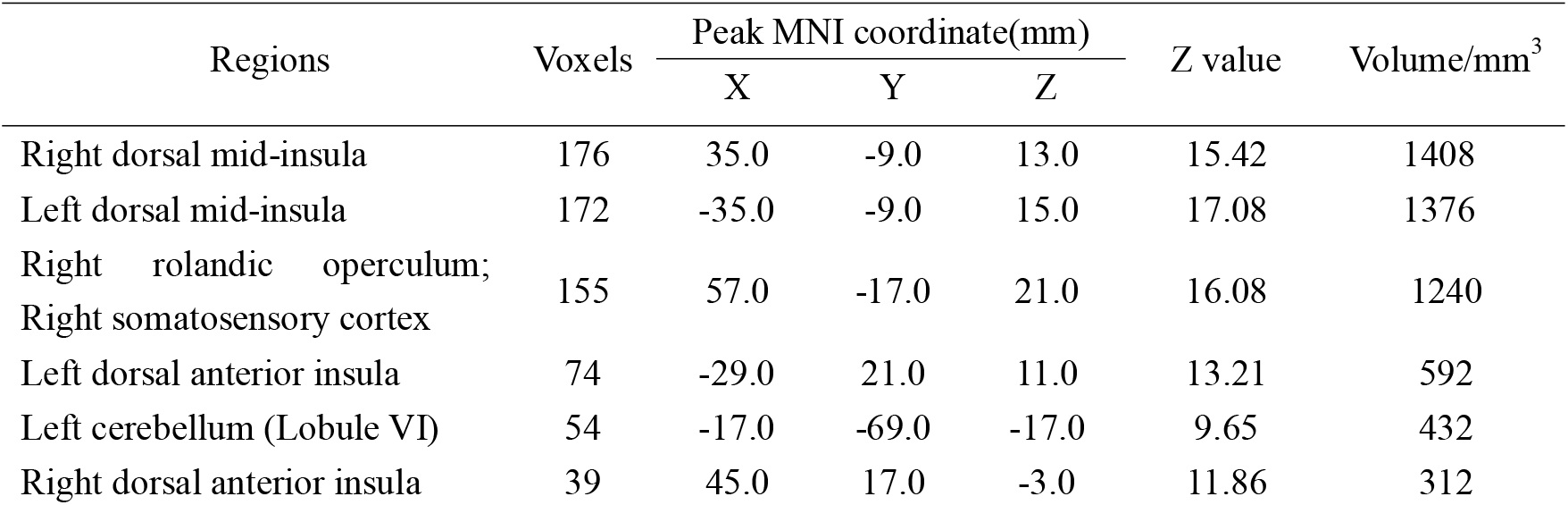
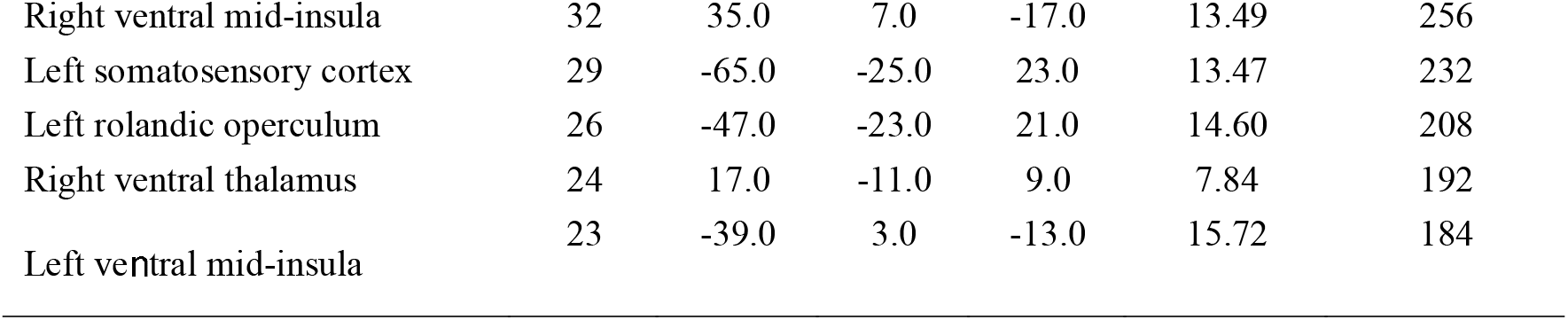
Gustatory activation areas responsive to taste.

**Figure 2.**
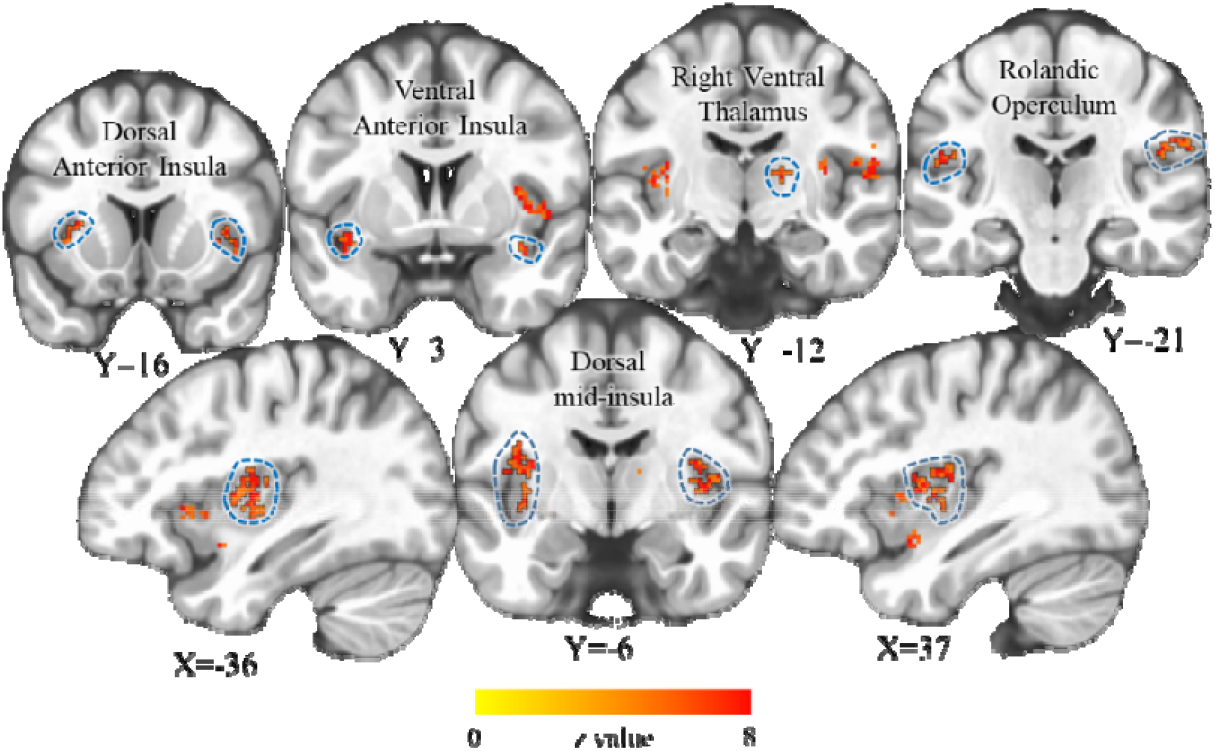
The gustatory activation areas of the brain. The gustatory cortex exhibited more significant BOLD responses to five basic tastes relative to tasteless included bilateral dorsal mid-insula, dorsal anterior insula, ventral anterior insula, rolandic operculum, and right ventral thalamus (P<0.00001, FDR corrected).

#### 3.2.2 Five taste responses maps in the insular cortex

At the subject level, the five taste responses maps of four subjects in the insular cortex were shown in Fig.3.In each subject, the five taste response maps showed an important, roughly the same bilateral dorsal mid-insula location in the central insular sulcus. Across four subjects, the five taste response maps demonstrated an interindividual variability of response intensities. The mean statistic contrast map of four subjects exhibited a significant activation of bilateral precentral and central insular sulcus(Naidich et al., 2004; Richardson & Fridriksson, 2016; Uddin, Nomi, Hebert-Seropian, Ghaziri, & Boucher, 2017).

**Figure 3.**
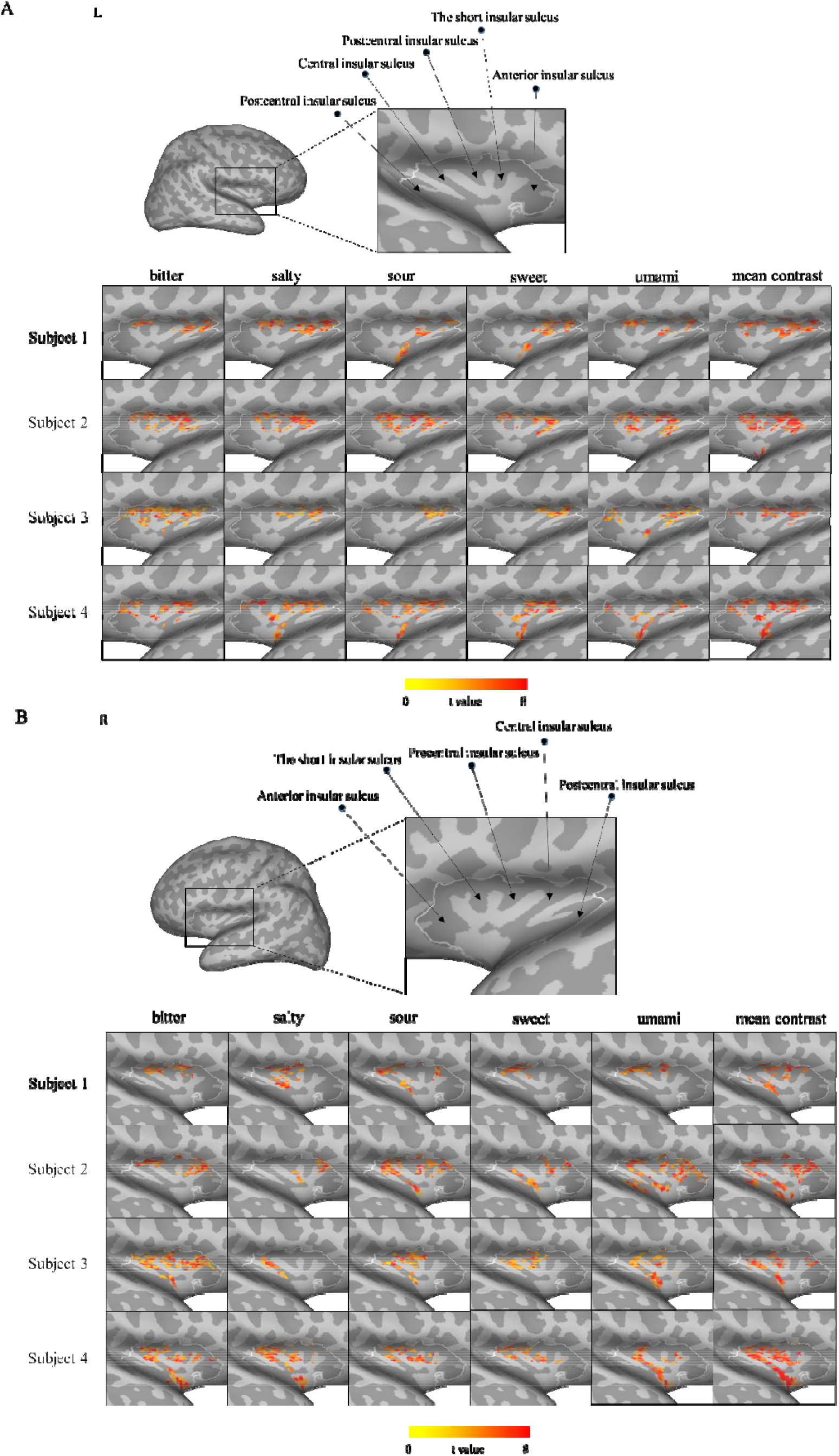
Activation maps for four subjects’ bitter, salty, sour, sweet, and umami stimuli. **A**. The left hemisphere of five taste responses map in four subjects. **B**. The right hemisphere of the five taste responses map in four subjects. L, left; R, right.

All tastes showed similar active areas at the group level, and three taste-responding clusters were observed at each hemisphere. Two of the three clusters were located in the dorsal mid-insula, with one in the mid of the dorsal mid-insula and the other closed to the parietal operculum. Another cluster was located in the dorsal left anterior insula (Fig. 4B).In addition, masks of taste-active voxels were generated for each taste. The masks were highly overlapped within the central and precentral insular sulcus (Fig. 4D), which meant the central sulcus was responsible for the recognition of five tastes. In addition, the activations in the central insular sulcus showed high symmetry and strong activation, suggesting taste is activated bilaterally.

**Figure 4.**
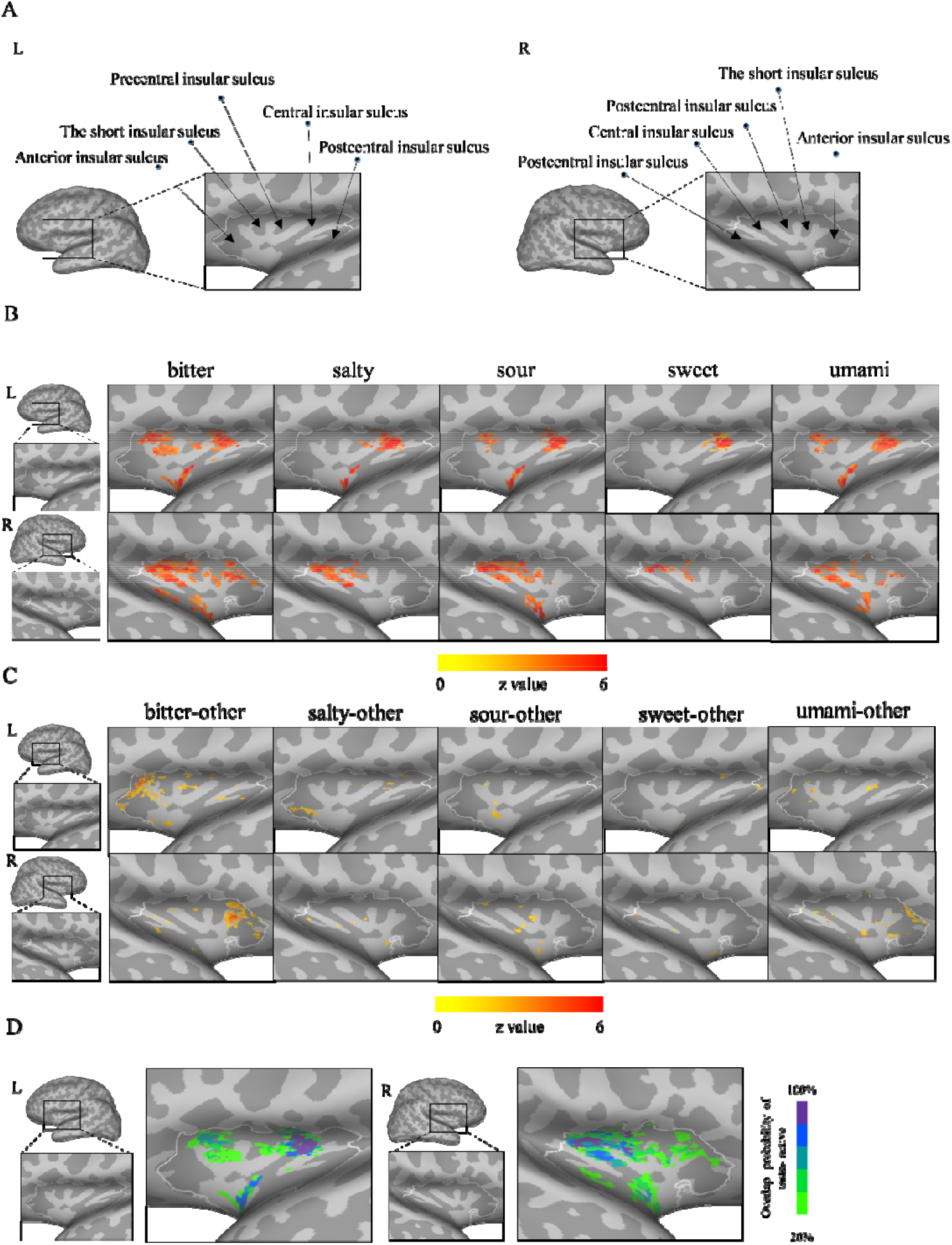
Group analysis results. **A**. The left and right hemisphere insula is shown with sulci labeled. **B**. The main three clusters are located in the dorsal middle insula, ventral middle insula, and dorsal anterior insula in the left hemisphere. The main three clusters are located in the parietal lobe-dorsal middle insula, mid-dorsal middle insula, and dorsal anterior insula in the right hemisphere. **C**. taste preference maps of insular cortex. Maps are thresholded at z = 1.96(P<0.05, cluster unclustered). Contrast analyses of taste preference areas demonstrate no clear evidence of topographical organization. **D**. The probability of overlapping of the taste active zone in the insula, where the five tastes were reliably activated (P<0.01, FDR corrected).L, left; R, right.

#### 3.2.3 Taste preference areas analysis

Contrasts between each taste and the average of the remaining tastes were calculated (bitter-other, salty-other, sour-other, sweet-other, umami-other) to identify areas within the insular cortex preferentially activated by either taste. Our voxel-wise false discovery rate corrected (P<0.05) results did not provide evidence of any regions of the insular cortex to be exhibiting a significant preference for either bitter, salty, sour, sweet or umami (Fig. 4B; uncorrected z maps for display only).

#### 3.2.4 MVPA results

We implemented multivariate pattern analysis using the 20 beta-coefficient maps of each subject within the dorsal mid-insula and anterior insula identified by mean taste contrast. The decoding results of every subject all failed to identify the five taste activity patterns. We were unable to distinguish the taste-selective activity patterns with an average classification accuracy of 20.18 ± 5.70% within the left dorsal mid-insula and 22.24 ± 4.97% within the right dorsal mid-insula (Fig.5A). The taste-selective activity patterns were also not distinguished with an average classification accuracy of 21.33 ± 2.98% within the left dorsal anterior insula and 20.48 ± 3.56% within the right dorsal anterior insula (Fig.5B). The MVPA classification results near the random level revealed the variability of taste-responsive patterns within different sessions.

**Figure 5.**
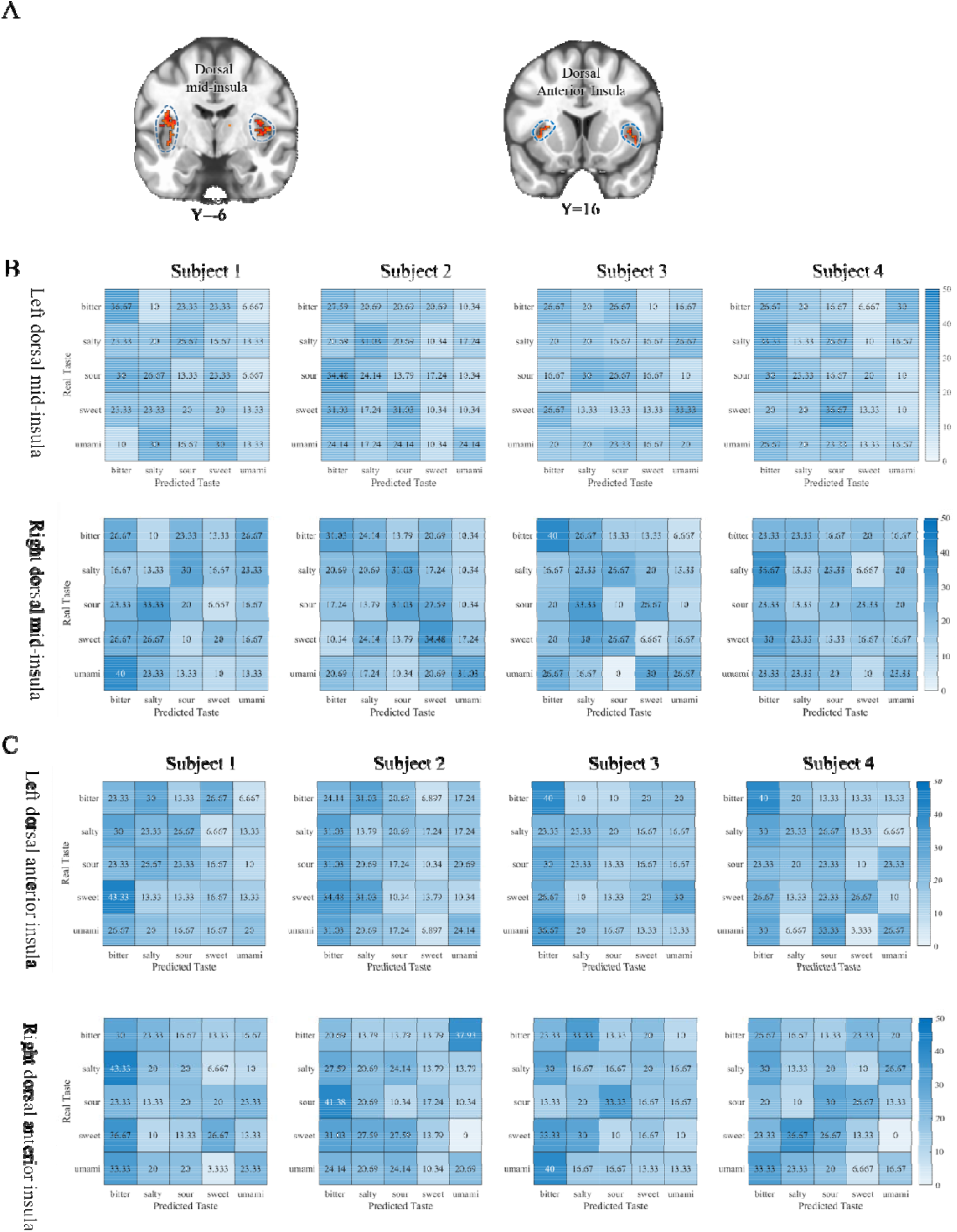
Multivariate pattern analyses results of region-of-interest within the bilateral dorsal mid-insula and dorsal anterior insula. **A**.ROIs of the dorsal mid-insula and the dorsal anterior insula.**B**. The confusion matrix of the dorsal anterior insula. **C**.The confusion matrix of the dorsal anterior insula.

**Figure 6.**
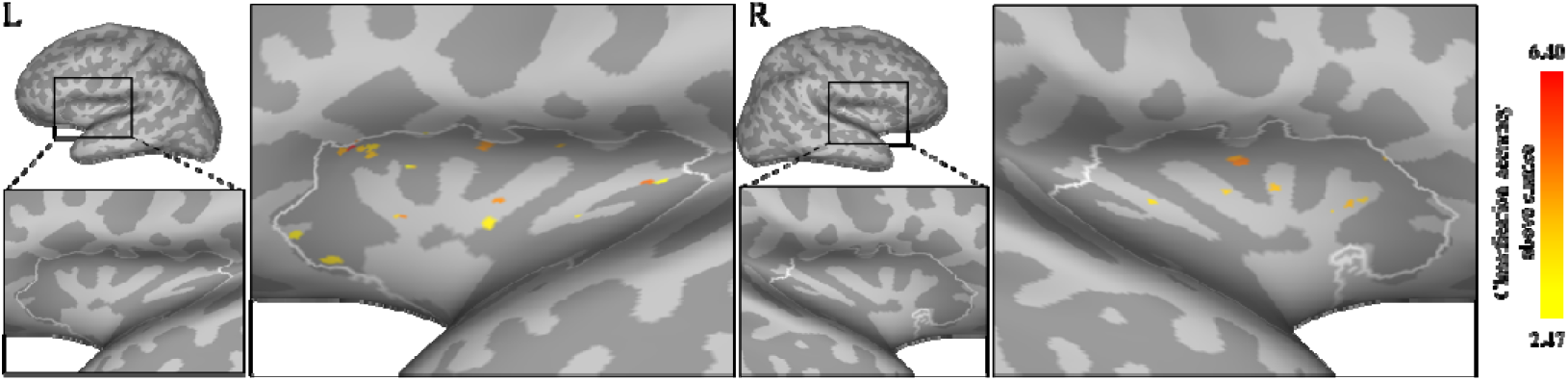
Multivariate searchlight analysis results within the insular cortex. The insular cortex shows little discrimination of taste activation patterns. The maps above show uncorrected results at a low statistical threshold (P<0.05, cluster unclustered).L, left; R, right.

Then we used the multivariate searchlight analysis to identify the regions which proved to have above-chance classification accuracy for discriminating between five tastes responses in the insular cortex. No voxel was found to survive after FDR multiple comparisons (P<0.5). Our multivariate searchlight analysis exhibited little discrimination of taste activation patterns in the insular cortex (Fig. 5; uncorrected maps presented for display only).

## 4 Discussion

This study aimed to derive comprehensive gustatory topologic maps of the human gustatory insular cortex. A self-developed taste delivery system was conducted to deliver the five basic taste and tasteless solutions to the subjects’ tongues, controlling for the effect of swallowing and the tastant temperature. Four participants were scanned over five sessions while performing the tasting task. Results show that the tastes activate a wide range of brain areas within the dorsal mid-insula, somatosensory cortex, dorsal anterior insula and ventral mid-insula. Both the individual-level and group-level univariate analysis demonstrates that each basic taste activates similar insular areas, which are mainly in the precentral and central insular sulcus. A further MVPA was unable to distinguish the basic tastes either in the dorsal anterior insula or the dorsal middle insula.

Another change of our experiment design compared to previous taste experiments is the continuously flowing liquids within the subjects’ mouth – taste (at task state) or tasteless (at baseline state) liquids were continuously pumped into the subjects’ mouth. Thus, the somatosensory responses to the liquid flows were minimized, and the bilateral gustatory insular cortex became the most activated area in the present work. We used sucralose rather than sucrose for the sweet solutions. Because sucrose solutions (for example, 0.6Mol/L sucrose) are so viscous that it is not fully washed out during the washing stage of the experiment, leading to constant sweetness feeling. Sucralose is about 600 times as sweetening compared to sucrose and activates the human insular cortex in a similar pattern but lower intensity compared to sucrose (Frank et al., 2008). The sucralose solutions (2mMol/L) used in the present work were easily washed out, and eliminate the taste interference.

Human gustatory experience is a complex process referring to the perception of taste quality, intensity, and pleasantness (Bender, Veldhuizen, Meltzer, Gitelman, & Small, 2009; Small, 2010). Taste receptor cells (TRCs) clustered in taste buds on the tongue are the first place where taste information processing begins. After taste information discharges from the TRCs, it flows along the chorda tympani (the three cranial nerves XII) and glossopharyngeal (cranial nerves IX), and lingual (cranial nerves X) to the nucleus of the solitary tract (NST). In human beings, taste signal is carried from the NST to the ventral posteromedial nucleus of the thalamus (VPMNT) and then to the gustatory insular cortex (Flynn, 1999). The right ventral thalamus cluster of the activation for five basic tastes further confirmed the experimental design was reliable in locating the taste functional regions of the brain.

Our observation of taste responses in the bilateral dorsal anterior insula, dorsal middle insula and parietal operculum is consistent with previous researches (Avery et al., 2020; Veldhuizen et al., 2011; Yeung, Goto, & Leung, 2017; Yeung et al., 2018). The dorsal mid-insula is thought to be the primary gustatory cortex (Bender et al., 2009; Small, 2010). The human secondary somatosensory cortex is located at the parietal operculum (Eickhoff, Schleicher, Zilles, & Amunts, 2006). The dorsal anterior insula was likely to be an integration area of multiple sensory information and also engaged in cognition and attention function (Kurth, Zilles, Fox, Laird, & Eickhoff, 2010). Our work furtherly shows that the taste response areas are within the central and precentral insular sulcus of the human middle insula.

However, the gustatory insular cortex responding to each basic taste is substantially overlapped. The taste preference areas analysis provides no evidence of any regions of the insular cortex exhibiting a significant preference for either bitter, salty, sour, sweet, or umami. The present work shows that the taste response areas are overlapped at both the individual level and the group level. A further MVPA was also unable to distinguish the basic tastes either in the dorsal anterior insula or the dorsal middle insula. fMRI MVPA fails when neuronal responses are weakly clustered(Dubois, de Berker, & Tsao, 2015). Thus, failure of the fMRI MVPA in the present study suggests the possibility of weakly clustered distribution of the taste-preference neurons in the human insular cortex. This is compatible with the most recent mammal studies, which have observed taste-preference neurons spatially distributing throughout the superficial layer of GC with no sign of spatial clustering or topography (K. Chen et al., 2021; Fletcher et al., 2017). A previous MVPA work distinguishes among the sweet, salty, and sour tastants above chance levels by the signal from the middle insula(Avery et al., 2020). There are two noticeable differences between Avery et al. (2020)’s and ours: First, the action of swallowing was avoided in the present study, and so was the swallowing related neural activity (S. Yang, T. J. Ross, Y. Zhang, E. A. Stein, & Y. Yang, 2005); Second, the duration of taste stimulus is much shorter in our work (2.6 seconds *vs*.20 seconds), and it lessens the potential effect of top-down affective responses or cognitive activities (Craig, 2002; Singer, Critchley, & Preuschoff, 2009) in the insula.

Although taste quality information is not represented by spatially unique clusters of neurons within the insular cortex, a microscale organization of taste selective responses neurons in the human insular cortex is still possible. In addition, specific temporal patterns of neuronal activity may represent both quality and valence. The temporal resolutions of fMRI may limit the study of the dynamic of taste perception, which could distinguish the taste quality information at the voxel level (Crouzet, Busch, & Ohla, 2015; Iannilli, Noennig, Hummel, & Schoenfeld, 2014). Imaging techniques with higher temporal and spatial resolution would be essential to explore the chemotopic organization of taste quality in the insular cortex.

## 5 Conclusions

The present study investigated the BOLD response of taste stimuli while controlling for effects induced by swallowing and tastant temperature. The results showed that the bilateral mid-insula activated most during the taste task, and the active areas were mainly in the precentral and central insular sulcus. But the areas responding to the five basic tastes were overlapped in the gustatory insular cortex. Furthermore, we did not identify any regions within the insula that exhibited a preference for a specific taste. Similarly, the multivariate pattern analysis results suggested it was difficult to catch the fixed pattern of five taste responses in the human insular cortex. In conclusion, our results support the idea that taste quality is not represented topographically in the human insula.

## Acknowledgments

This work was supported by the National Natural Science Foundation of China (grant nos. 81701665, 21876041) and the support of the Foundation Students’ Innovation and Entrepreneurship Foundation of USTC (nos.CY2022X09).

## Data and code availability statement

The data and code used in this paper are available via a request to the corresponding author.

